# Biomechanical and biochemical assessment of YB-1 expression in melanoma cells

**DOI:** 10.1101/2021.12.29.474412

**Authors:** Anna Cykowska, Ulf Krister Hofmann, Aadhya Tiwari, Corinna Kosnopfel, Rosa Riester, Marina Danalache

## Abstract

Malignant melanoma is the most lethal form of skin cancer; its incidence has increased over the last five decades. Y-box binding protein 1 (YB-1) plays a prominent role in mediating metastatic behavior by promoting epithelial-to-mesenchymal transition (EMT) processes. Migratory melanoma cells exhibit two major phenotypes: elongated mesenchymal or rounded amoeboid. The actomyosin cytoskeleton is key in both phenotypes, but intermediate filaments also undergo a significant rearrangement process, switching from cytokeratin-rich to vimentin and nestin-rich network. In this study, we aimed to investigate to what extent YB-1 impacts the biomechanical (cell stiffness) and biochemical aspects of melanoma cells and their cytoskeleton. To this end, we subjected A375 YB-1 knock-out and parental cells to atomic force microscopy investigations (stiffness determination), immunolabelling, and proteome analysis. We found that YB-1 expressing cells were significantly stiffer compared to the corresponding YB-1 knock-out cell line. Proteomic analysis revealed that expression of YB-1 results in a strong co-expression of nestin, vimentin, fascin-1, and septin-9. In the YB-1 knock-out nestin was completely depleted, but zyxin was strongly upregulated. Collectively, our results showed that YB-1 knock-out acquires some characteristics of mesenchymal phenotype but lacks important markers of malignancy and invasiveness such as nestin or vimentin. We posit that there is an association of YB-1 expression with an amoeboid phenotype, which would explain the increased migratory capacity.

## 1. Introduction

Malignant melanoma is among the most lethal form of skin cancer and the prognosis for affected patients is grim, with an estimated 5-years survival rate ranging between 5-19% (1).

Y-box binding protein 1 (YB-1) is a multifunctional member of the cold-shock protein superfamily that has a dual role as a transcription factor in the nucleus as well as a translational regulator in the cytoplasm (2,3). It is also a key player in neoplastic development and progression as upregulation and nuclear translocation of YB-1 is linked to patient relapse and poor prognosis in a plethora of cancer types (4-6). YB-1 also plays an important role in mediating metastatic behavior (7) and it is a biomarker of exacerbated tumor progression (8). Kosnopfel et al. showed that the cytoplasmic activity of YB-1 triggers tumorigenicity and metastatic potential of melanoma cells by promoting epithelial-to-mesenchymal transition (EMT) processes (9).

The metastatic behavior of malignant melanoma depends on its ability to migrate and invade. Migratory melanoma cells exhibit two major phenotypes: elongated mesenchymal or rounded amoeboid. Mesenchymal migration is a protrusion-dependent mode mediated by polarized signaling of GTPases Rac1 and Cdc42 and is characterized by focal adhesion formation and lower actomyosin contractility (10). Rounded amoeboid invasion uses unstable blebs that can squeeze through the matrix, therefore exhibits a low degree of β1integrin-mediated adhesion, reduced focal adhesion size, high actomyosin contractility, blebs formation, and high cytokine signaling (11).

Even though both migratory strategies are the extreme spectrum, the actomyosin cytoskeleton is key in both phenotypes; the contractile cortex is important for amoeboid-rounded to intermediate forms (12) but some degree of contractility is also required to retract protrusions in elongated-mesenchymal phenotype (13).

From a structural perspective, the cytoskeleton comprises three main components: the actin cytoskeleton, the microtubule network, and the intermediate filaments. Although these components act synergistically, the driving force required for cell migration is the actin cytoskeleton (14). Alongside the actin cytoskeleton, microtubules also regulate the metastatic potential via the crosstalk with actin (15).

Coexisting with actin and microtubules, intermediate filaments are responsible for traction forces between cells and protect the cells from disruptions. Throughout EMT, intermediate filaments undergo a significant rearrangement process, switching from cytokeratin-rich to vimentin and nestin-rich networks, which entails an enhanced cell motility capacity (16).

YB-1 expression is upregulated during melanoma progression and its activity is regulated by the constitutively active MAPK signaling pathway as addressed in (9,17,18) and are not further elucidated in this work. Along with the promotion of EMT-like processes by cytoplasmic YB-1, Kosnopfel et al. previously showed that YB-1 is secreted from melanoma cells in a progression-stage dependent manner and stimulates melanoma cell migration and invasion (19). This data provides a sound rationale for the analyses conducted in this study.

In this study, we aimed to investigate to what extent YB-1 impacts the biochemical properties of melanoma cells and their cytoskeleton. In particular, we investigated the effect YB-1 presence or absence has on cellular elastic features at a nanoscale. We further analyzed its function both quantitatively and qualitatively by means of proteomics and immunohistochemical labeling.

## 2. Materials & Methods

### 2.1. Cell lines and culture

A375 and MelJuso cells (ATCC, Manassas, VA, USA and DSMZ, Braunschweig, Germany) were maintained in RPMI-1640 with L-glutamine (Gibco, Life Technologies, Darmstadt, Germany) supplemented with 10% (v/v) fetal calf serum (Merck Biochrom, Berlin, Germany) and 1% (v/v) penicillin/streptomycin (Thermo Fisher Scientific, Waltham, MA, USA) in an incubator at 37°C as previously described (19).

For all experiments, cells with passage numbers of 5–20 at about 80% confluency were harvested. All melanoma cell lines were used for no longer than 2 months after thawing from the frozen stock.

### 2.2. YB-1 knock-out in A375 cells

YB-1 gene knock-out (CRISPR YB-1^KO^) for the A375 cell line was conducted using CRISPR/Cas9 mediated genome engineering and single-cell clones with effective YB-1 knock-out identified as described previously (9,19). The sequences of the sgRNAs used for the YB-1 knock-out along with the lentiviral transfer vector (lentiCRISPRv2) used for the transduction of the melanoma were also described previously (18).

### 2.3. Biomechanical characterization by Atomic Force Microscopy (AFM)

To measure the stiffness of cells, we used an atomic force microscope (AFM) system (CellHesion 200, JPK Instruments, Berlin, Germany) mounted onto an inverted light microscope (AxioObserver D1, Carl Zeiss Microscopy, Jena, Germany), which allowed us to optically align the cantilever to the region of interest and to simultaneously measure specific regions of interest on individual cells (Figure 1). Calibration of the cantilever was done on a surface of an empty petri dish filled with a Leibovitz’s L-15 medium without L-glutamine (Merck KGaA, Darmstadt, Germany) pre-warmed to 37°C. It was performed on the retracted curve and the spring constant was determined by using the thermal noise method incorporated into the device software (JPK Instruments, Berlin, Germany). Measurements were performed in force spectroscopy mode by recording single force-distance curves at the position of interest without laterally scanning the sample. The indentation was performed using a spherical tip of 5 µm (Model: SAA-SPH-5UM, k=0.2 N/m, Bruker, Billerica, MA, USA). Cells were grown for 48 hours prior to the AFM biomechanical measurements on the bottom of the dishes (TPP Techno Plastic Products AG, Trasadingen, Switzerland) until sufficient confluency was reached (± 80%). The culture medium (described in 2.1) was changed 24 and 48 hours after seeding. Prior to mechanical measurements, the culture medium was removed, and the cells were covered with Leibovitz’s L-15 medium without L-glutamine (Merck KGaA). To eliminate the confounding effects of neighboring cells on cytoskeleton arrangement and morphology, randomly selected single cells were measured (Figure 1). To evaluate the elastic properties of A375 and MelJuso cells, we indented the selected cells approximately over the perinuclear region of individual cells. We measured 30 cells per condition and each cell was automatically measured five times at a single point. Indentation curves were sampled at 2 kHz, with a force trigger of ∼10 nN and a velocity of 5 µm/sec. The Young’s modulus for each individual cell was calculated from the force-distance curves from each measurement by using the Hertz fit model incorporated in the data processing software (JPK Instruments), with the Poisson’s ratio set to 0.5.

**Figure 1.**
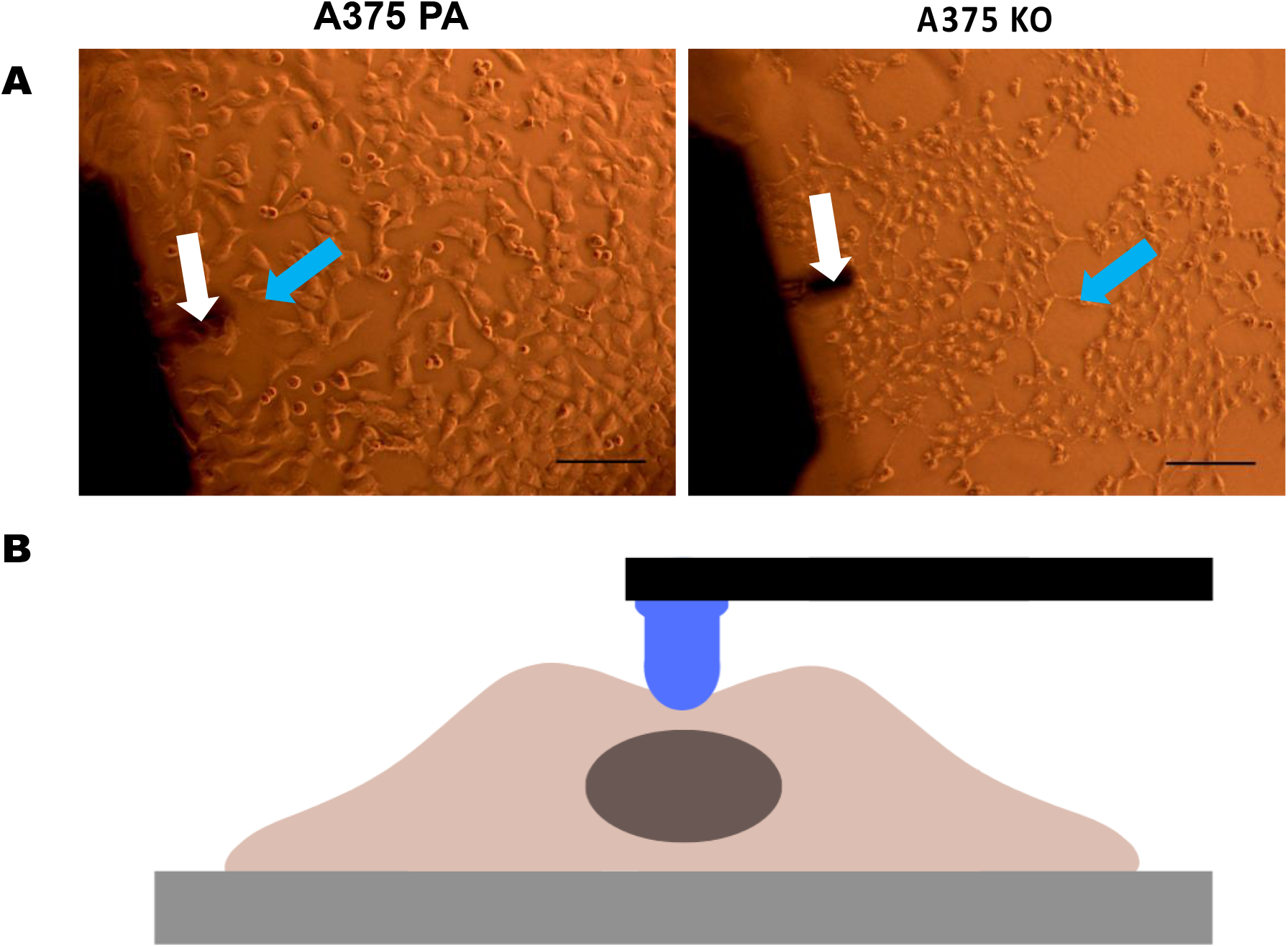
Microscopic images representing the regions of interest subjected to AFM measurements. (**A**) Cell lines A375 parental and A375 YB-1 knock-out as well as MelJuso parental were subjected to AFM measurements. White arrows indicate the cantilever employed for indentations. Blue arrows show the region of interest in an exemplary cell. (**B**) Schematic drawing of a single cell indentation by a cantilever. Images were acquired with the inverted AxioObserver D1 light microscope attached to the AFM system at a 10× magnification. Scale bar (black) represents 30 µm. Abbreviations: AFM -atomic force microscopy, PA -parental, KO - knock-out.

Since the stiffness of tumor cells can be an indicator of metastatic potential (20), alongside the A375 cells, MelJuso YB-1 expressing cells were also subjected to cellular stiffness measurements to investigate the possible heterogeneity of elastic features that may vary within cancerous tumors.

### 2.4. NanoLC-MS/MS analysis and data processing

Prior to the proteome analysis, YB-1 presence/absence in cells lines was verified via Western blot analysis as previously described (9). For the proteomic investigations, protein extracts were purified using SDS PAGE (Thermo Scientific). Coomassie-stained gel pieces were excised and in-gel digested using trypsin as described previously (21). After desalting using C18 Stage tips extracted peptides were separated on an Easy-nLC 1200 system coupled to a Q Exactive HF mass spectrometer (both Thermo Scientific) as described elsewhere by Kliza et al (22) with slight modifications. The peptide mixtures were separated using a 1 hour gradient. The seven most intense precursor ions were sequentially fragmented in each scan cycle using higher-energy collisional dissociation (HCD) fragmentation. Acquired MS spectra were processed with MaxQuant software package version 1.6.7.0 with an integrated Andromeda search engine (23). Database search was performed against a target-decoy *Homo sapiens* database obtained from Uniprot, containing 96,817 protein entries and 286 commonly observed contaminants. Endoprotease trypsin was defined as a protease with a maximum of two missed cleavages. Oxidation of methionine and N-terminal acetylation were specified as variable modifications, whereas carbamidomethylation on cysteine was set as fixed modification. The initial maximum allowed mass tolerance was set to 4.5 parts per million (ppm) for precursor ions and 20 ppm for fragment ions. The LFQ (Label-FreeQuantification) algorithm was enabled, as well as matches between runs (24) and LFQ protein intensities were used for relative protein quantification of three replicates. Gene Ontology (GO) Enrichment Analysis of biological processes and molecular function was conducted using ShinyGO v0.741 (25) (http://bioinformatics.sdstate.edu/go/) with Fisher’s exact test, false discovery rate (FDR) correction, and selecting a 0.05 p-value cut-off. ShinyGO was also used to generate the lollipop charts and the relationship networks between the enriched pathways.

### 2.5. Immunofluorescence

Cells were washed 3 times with phosphate-buffered saline (PBS) and fixed with 2% (w/v) formaldehyde and PBS for 15 mins. The chambers were washed three times for 5 minutes in PBS. Subsequently, the permeabilization was done with 0.2 % Triton X-100, PBS, and 1% BSA and the blocking with a mix of 3% (w/v) bovine serum albumin and PBS as a blocking agent for 30 min. This process was followed by incubation with collagen I (mouse, 1:100, #ab90395, abcam, Cambridge, UK), collagen II (mouse, 1:100, # MA-12789, Thermo Scientific, Waltham, MA, USA), collagen III (rabbit, 1:100, #ab7778, abcam), beta-tubulin (rabbit, 1:100 #2129 9F3, Cell Signaling, Danvers, MA, USA), fibrillin I (mouse, 1:100, #MA512770, Thermo Scientific), YB-1 (rabbit, 1:100, #ab76148, abcam), beta actin (1:100, #ab8227, abcam), vimentin (1:100, Rabbit #D21H3, Cell Signaling), and nestin (1:50, Mouse #sc23927, Santa Cruz, Dallas, TX, USA) primary antibodies as well as F-actin (1:100, Alexa Fluor 596 phalloidin #A12380, Thermo Scientific), G-actin staining (DNase I Alexa Fluor 488 conjugate, #D12371 Molecular Probes, Eugene, OR, USA) conjugated antibodies in 2.5% (w/v) bovine serum albumin in PBS overnight at 4°C in a humidity chamber. Afterward, sections with primary antibodies were incubated with secondary antibodies (Alexa Fluor-594 goat anti-rabbit IgG, #a21429, Thermo Scientific; Alexa Fluor 488 goat anti-mouse IgG, #ab-150116, abcam; Alexa Fluor 594 goat anti-mouse IgG, #a-21429, Thermo Scientific; Alexa Fluor 488 goat anti-rabbit IgG, #ab150116, abcam) for 2 h with a dilution of 1:100 in 0.5% (w/v) bovine serum albumin at room temperature in the dark. The chambers were washed three times for 5 minutes each in PBS. Nuclear staining was performed with 1% (v/v) DAPI (Life Technologies, Darmstadt, Germany) 5 min prior to imaging. Fluorescence-stained tissue sections were visualized with a Carl Zeiss Observer Z1 fluorescence microscope (Carl Zeiss Microscopy, Jena, Germany). All stainings were repeated twice. For negative controls, we omitted primary antibodies.

### 2.6. Statistical analysis

For further analyses, the arithmetic mean of the five AFM measurements per cell was used. Based on the non-normality of these means, AFM values are presented as median (minimum-maximum) and graphically displayed as boxplots. Descriptive statistics including standard deviation, and standard error of the mean of the AFM data are additionally displayed (Figure 2 B and Supplementary Figure 1). Inferential statistics were performed with the non-parametric Mann– Whitney U test to compare the differences between groups based on an alpha of 0.05. Statistical analysis was performed using the SPSS Statistics 22 (IBM Corp., Armonk, NY, USA) software. For the proteomics data, further *in-silico* analysis was conducted where data handling, statistical tests, and figure generation were performed using ProVision (https://provision.shinyapps.io/provision/) - an online open-source data analysis platform that allows for the analysis of MaxQuant files (26). Briefly, reverse database hits, contaminants, and proteins only identified by site modifications were removed. The file was further filtered for each protein group to contain at least two unique peptides. The assigned LFQ intensity values were subsequently log_2_ transformed to gain a normal distribution and further filtered for two values in at least one group. This resulted in a high confidence expression dataset and missing values were imputed from a truncated normal distribution of transformed LFQ intensities. Quantile plots were done within the ProVision application to check for data normality prior to statistical testing. Multiple hypothesis testing was corrected using the Benjamini-Hochberg FDR set at 0.05, and a two-fold cut-off was implemented.

**Figure 2.**
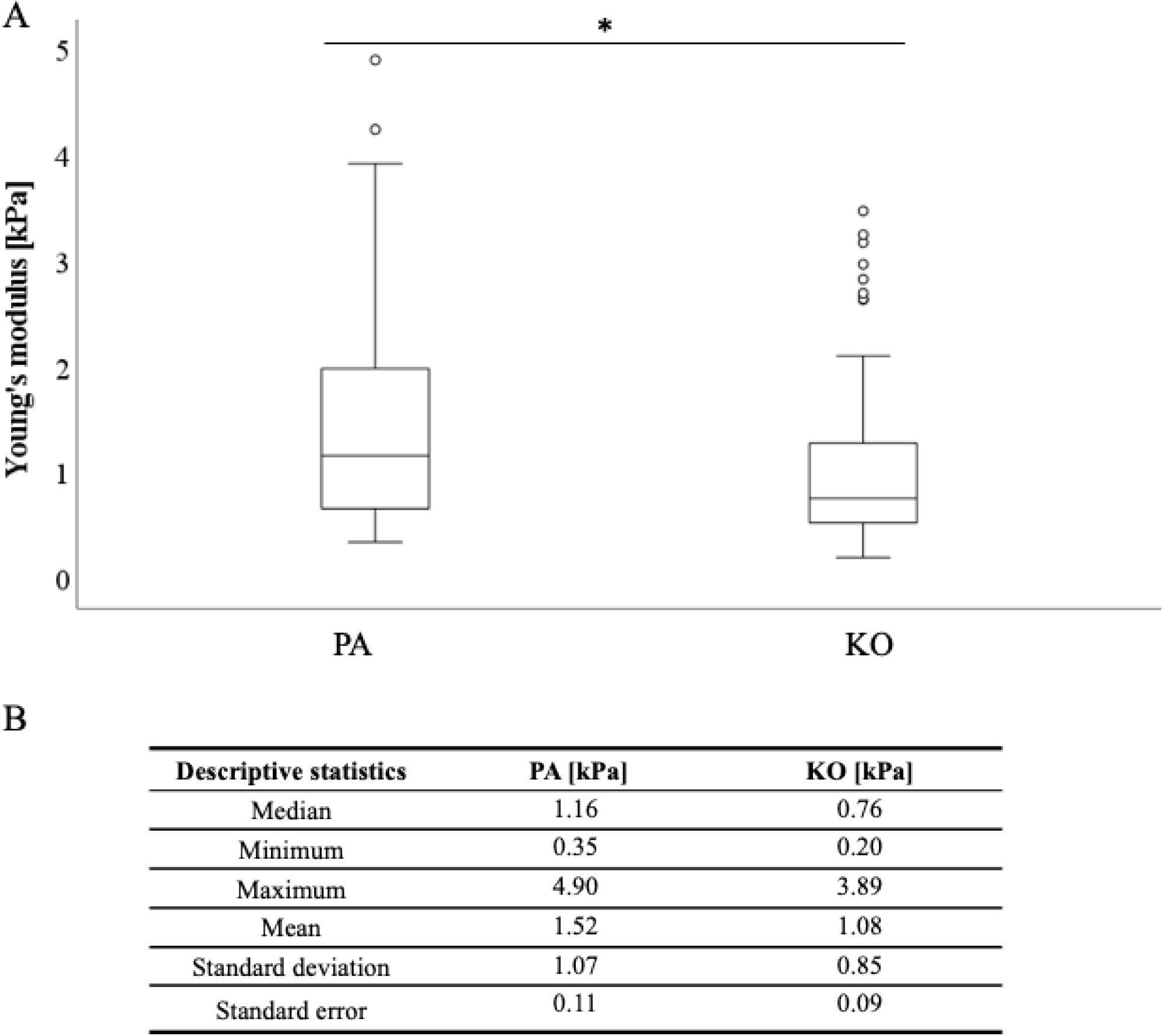
Cell stiffness as measured by Young’s modulus (kPa) is higher in the YB-1 expressing parental A375 cells when compared to the YB-1 knock-out cell line. Boxplots represent the difference in stiffness (kPa) measured by AFM between the two conditions (parental and YB-1 knock-out) (**A**), with the descriptive statistical analysis below (**B**). A significantly higher stiffness was observed in the A375 parental cell line compared to its corresponding YB-1 knock-out condition (p<0.001). *p < 0.05; Abbreviations: PA - parental; KO - knock-out; AFM - atomic force microscopy.

The protein-protein interaction analyses and enrichment analysis were performed using the STRING database (https://string-db.org/) by submitting the proteins with ranks to normal geneset analysis and performing the Functional Enrichment Analysis with the standard settings (medium confidence 0.4). The interaction network was important and visualized in the open-source Cytoscape 3.9.0 (National Institute of General Medical Sciences of the National Institutes of Health, USA, https://cytoscape.org) via Cytoscape StringApp and STRING Enrichment as previously described by Shannon et al (27).

## 3. Results

### 3.1. YB-1 renders cells stiffer

A total of 30 cells per cell line/condition were subjected to stiffness measurements in the perinuclear region via AFM, and each individual cell was automatically measured five times in three independent repetitions. Thus, a total of 1350 measurements were conducted (for each cell line/condition a total of 450 measurements). The measured cellular elastic moduli are depicted in Figure 2 for A375 cell line (YB-1 knock-out and parental) and in Supplementary Figure 1 for A375 – MelJuso comparison. Stiffness for the A375 parental cell line with endogenous YB-1 expression was significantly higher (p<0.001) than for the A375 YB-1 knock-out (Figure 2). To account for the possible heterogeneity of mechanical properties in tumor cells, the MelJuso cell line was concomitantly subjected to AFM stiffness measurements. A significant stiffness decrease was noted when comparing the two YB-1 expressing cell lines: A375 and MelJuso (p<0.001, Supplementary Figure 1). This complements the view of cancer as a heterogeneous pathology, showing the mechanical properties between cancerous tumors. Overall, absolute values were thereby reduced by 34% with a median of 1.16 kPa for the A375 parental to a median of 0.76 kPa for the A375 knock-out, and 27% for the MelJuso parental cell line (median of 0.84 kPa) respectively. We also observed different phenotypes in A375 parental and knock-out (Figure 1) where parent cells were more rounded and bigger while knock-out cells were smaller and more elongated.

### 3.2. Proteomic and STRING Analysis of Melanoma Response to YB-1 knock-out

The proteomic analysis aimed to further identify the main proteins and protein networks associated with the expression of YB-1 and their relation to the biomechanical and biochemical aspects of melanoma cells with a particular focus on cytoskeleton proteins.

A total of 136 significantly differentially expressed proteins (DEPs) were revealed of which 55 upregulated in the A375 parental cell line and 81 in the A375 YB-1 knock-out (Figure 3 A, B). The top 10 DEPs in each cell line were further visualized in the volcano plot (Figure 3 A) and the heatmap (Figure 4). Confirming our study design YB-1 was expressed in the parental cell line and was completely absent in the knock-out. Interestingly, nestin expressed the same pattern as YB-1: highly expressed in the A375 parental cell line and absent from the YB-1 knock-out cell line (Figure 3 A).

**Figure 3.**
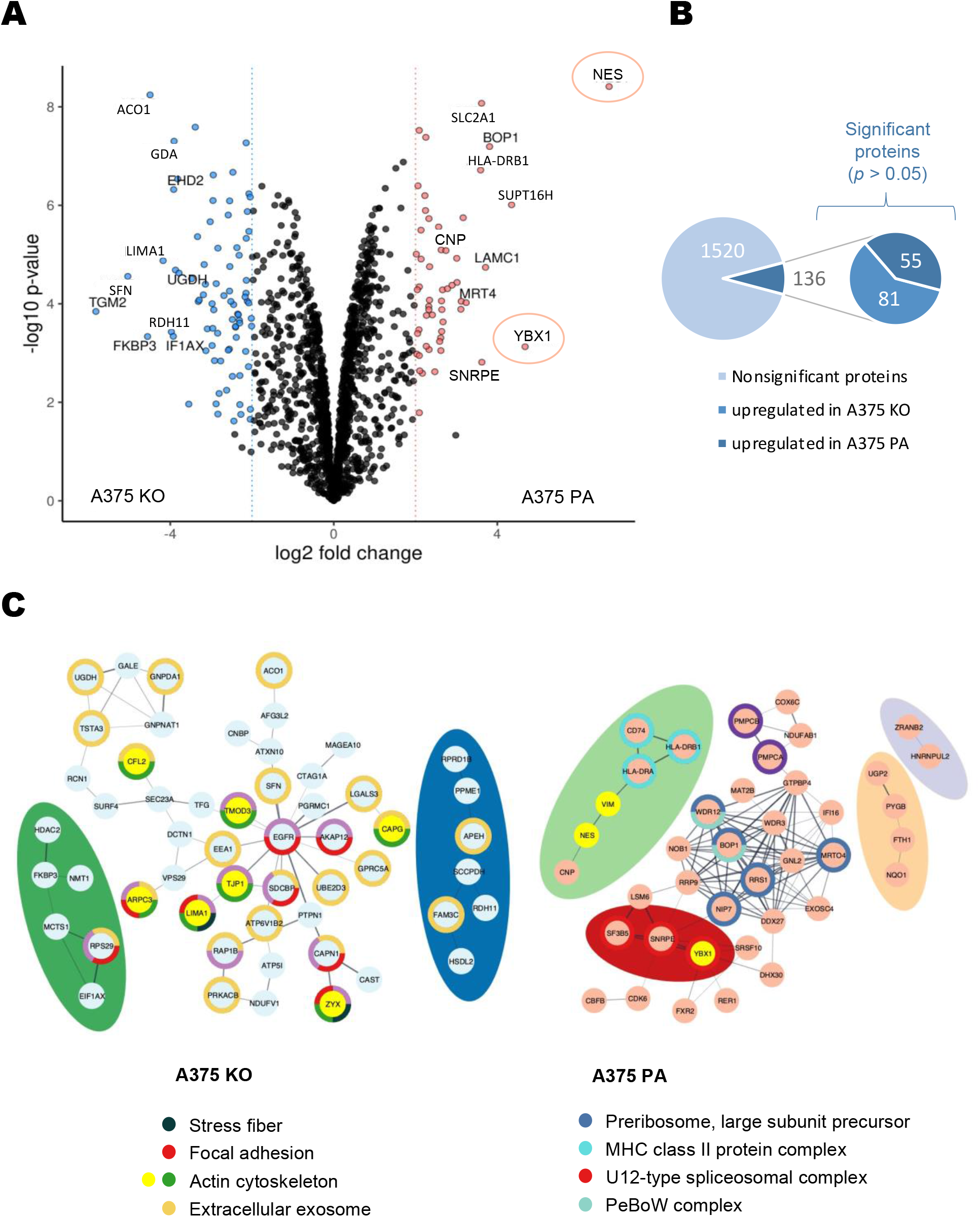
Analysis of significantly differentially expressed proteins (DEPs). **(A)** Volcano plot with the top 10 significant DEPs labeled in A375 parental and A375 YB-1 knock-out. Each dot represents a different protein. A p-value <0.05 was considered statistically significant, whereas a fold-change of 2 was set as the threshold (represented by vertical lines). Nestin and YB-1 were marked by a red circle. **(B)** A number of significantly regulated proteins compared with the total number of quantified proteins. Data revealed a total of 136 DEPs, including 55 significantly preferentially upregulated proteins in the A375 parental and 81 in the A375 YB-1 knock-out. **(C)** A STRING analysis showing protein-protein interaction networks and donut annotation of the top significantly enriched cellular components (gene ontology). Networks represent significantly differentially expressed proteins in A375 parental and A375 YB-1 knock-out cell lines. The disconnected nodes were removed. Only the top 5 cellular components are shown. In the A375 YB-1 knock-out cell line, the nodes in yellow indicate actin cytoskeleton. In A375 parental, the nodes in yellow indicate nestin, vimentin, and YB-1. Abbreviations: DEPs - differentially expressed proteins; PA - parental; KO - knock-out.

**Figure 4.**
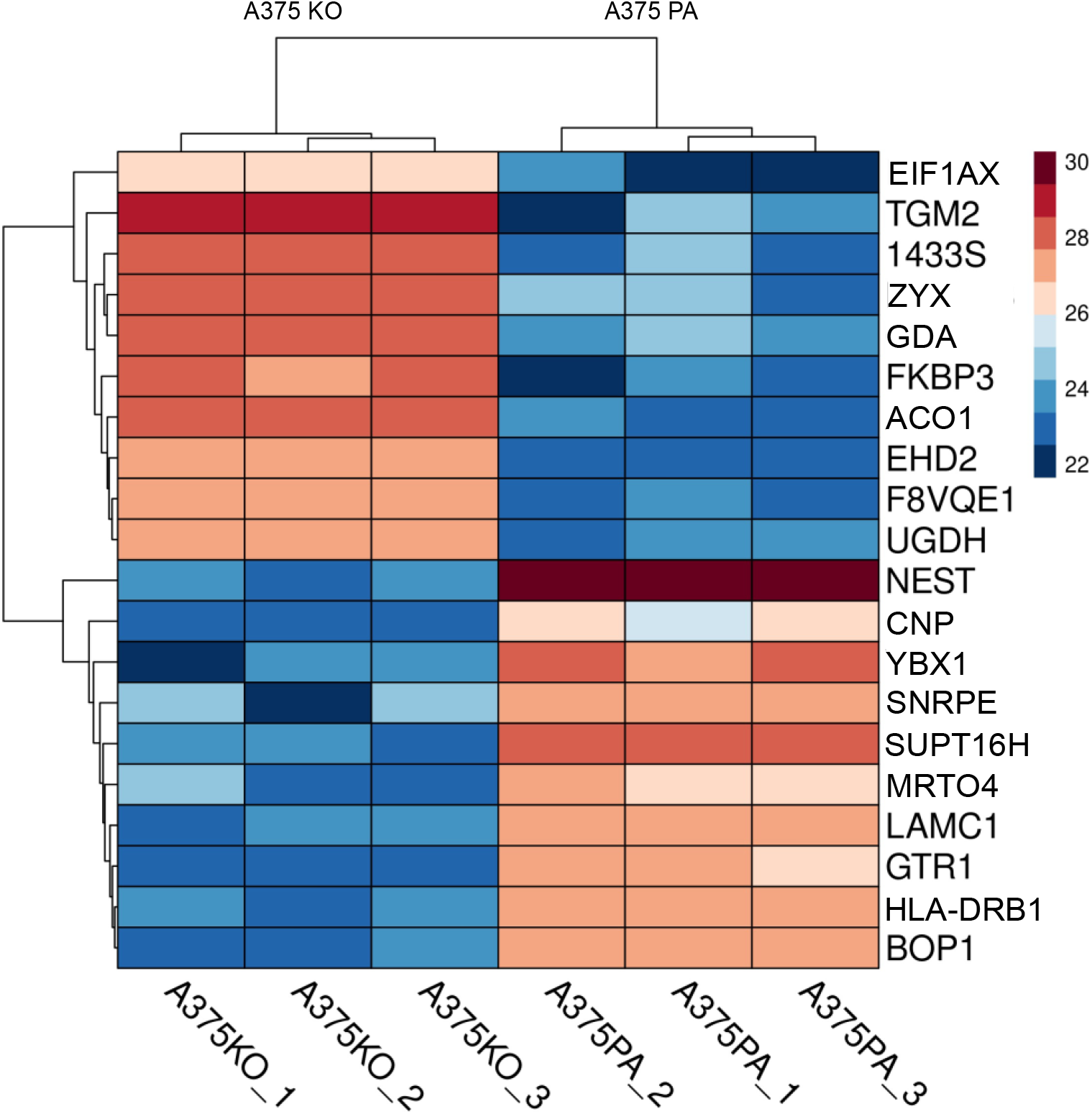
Heatmap of the Label-Free Quantification (LFQ) intensity values in triplicates in A375 parental and A375 YB-1 knock-out cells. The three vertical columns represent replicates of one cell line each and the LFQ intensity for each cell line is visualized. The top 10 significantly differentially expressed proteins (DEPs) for each cell line are shown. The color range of LFQ intensity extends from 30 (dark red) to 22 (dark blue). Analyzed and visualized in ProVision. Abbreviations: LFQ – Label-Free Quantification; PA - parental; KO – knock-out.

To shed light on the protein interactions occurring in our topmost cytoskeleton hits, we used a STRING approach to link and connect the information on known and predicted direct physical as well as indirect functional protein-protein interactions. Both cell lines showed distinct clusters with shared cellular functions (Figure 3 C). In the parental A375 cell line the most enriched cellular components were associated with preribosomes, MHC class II complex, spliceosomal and PeBoW complex. In contrast, in A375 YB-1 knock-out the most enriched cellular components were strongly related to the actin cytoskeleton, focal adhesion, stress fibers, and extracellular exosome (Figure 3 C).

To further elucidate the relationships between DEPs and gene functions, we employed enrichment analysis using gene ontology. The top enriched biological processes in both cell lines were sorted by the fold enrichment (defined as the percentage of genes in that list belonging to a pathway, divided by the corresponding percentage in the background) (Figure 5 A). The analysis revealed that in A375 parental cell line almost all overrepresented processes are related to RNA processing, transcription, and maturation. In the A375 YB-1 knock-out cell line, catabolic processes related to glucosamines were the most strongly overrepresented (Figure 5 B).

**Figure 5.**
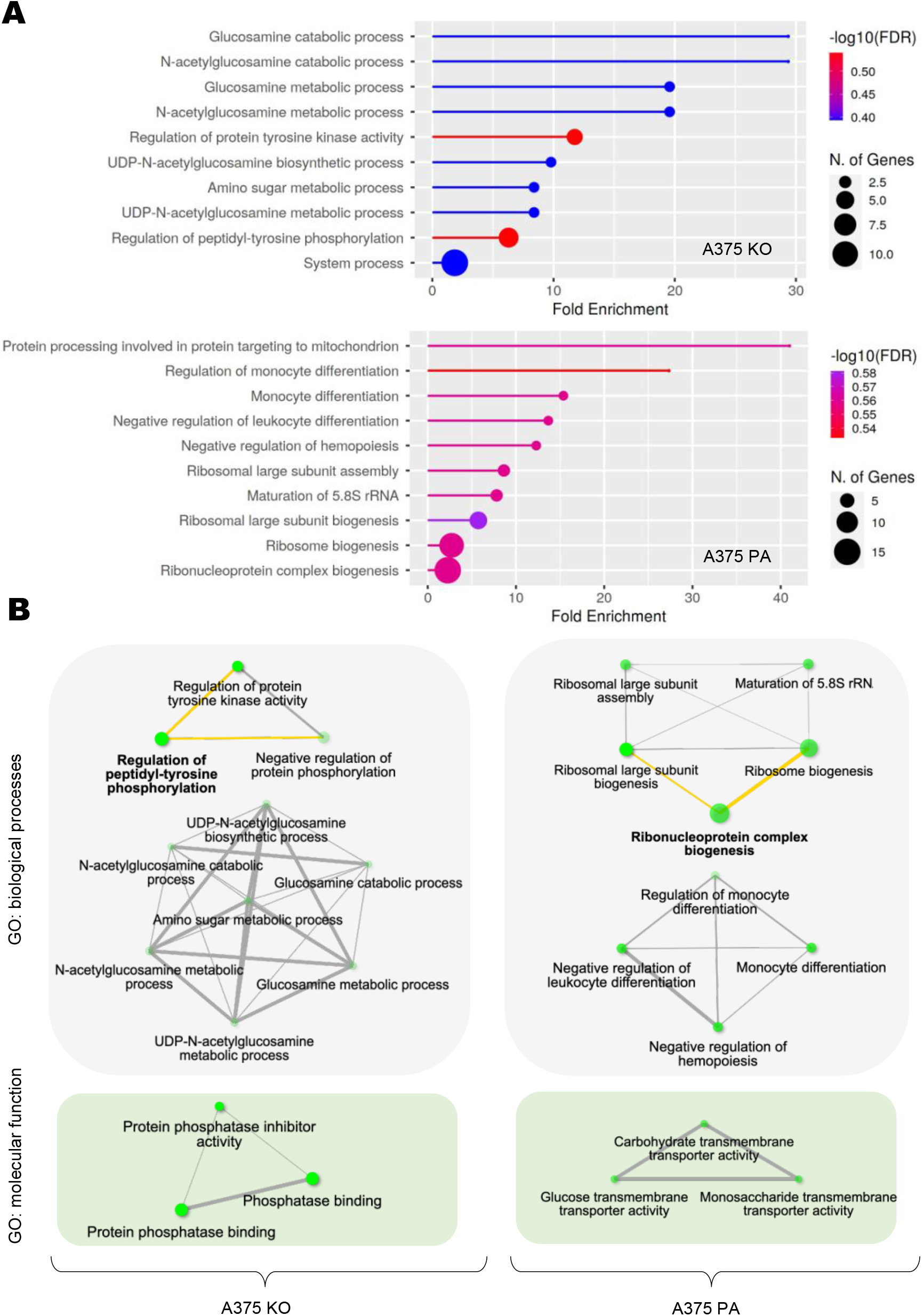
Gene Ontology (GO) Enrichment Analysis of biological processes and molecular function in the A375 parental and A375 YB-1 knock-out cell lines. (**A**) GO: biological processes enrichment analysis of the two cell lines. Lollipop charts of A375 parental and A375 YB-1 knock-out show the top 10 significantly enriched biological processes in each cell line sorted by fold enrichment. Analyzed and visualized using ShinyGO. (**B**) The relationship between enriched pathways in A375 parental and A375 YB-1 knock-out of GO: biological processes and GO: molecular pathways. Two nodes are connected if they share 20% (default) or more genes. Abbreviations: GO – Gene Ontology, PA – parental; KO – knock-out.

### 3.3. Qualitative assessment and localization of cytoskeleton proteins expression

To investigate the relationship between the biochemical and biomechanical changes as well as the cellular localization of selected proteins in YB-1 expressing and non-YB-1 expressing cell lines, we used immunofluorescence (IF) to determine the expression of the following proteins in the cultured cell lines: F-actin (filamentous), G-actin (globular), nestin, vimentin, collagen II, and beta-tubulin (Figure 6). Our selected targets corresponded to the cytoskeleton proteins of interest based on both the current literature and the results of the proteome analysis. Based on our immunolabelling results, G-actin is preferentially expressed in the nucleus or perinuclear region in the YB-1 expressing cell line (A375 parental, Figure 6 A). This is in contrast to the YB-1 knock-out cell line, where unpolymerized G-actin is widely expressed outside the nuclear region and intercellular contacts seem to follow precisely the contours of the cell (Figure 6 A). In contrast, in the parental cell line, F-actin clearly delineates cell boundary (Figure 6 B). The distribution of nestin and vimentin is also markedly different in the two cell types: in the A375 parental cell line, nestin is strongly expressed and concentrated in the perinuclear area (Figure 6 C) while in the A375 YB-1 knock-out cell nestin is barely visible. Vimentin on the other hand appears to produce a very strong signal and is more evenly distributed in the parental cell line within the cytoplasm (Figure 6 C). In YB-1 knock-out cell line, vimentin appears to be concentrated around the nucleus. Beta-tubulin network is more prominent and more distributed in the parental cell line compared to YB-1 knock-out (Figure 6 D).

**Figure 6.**
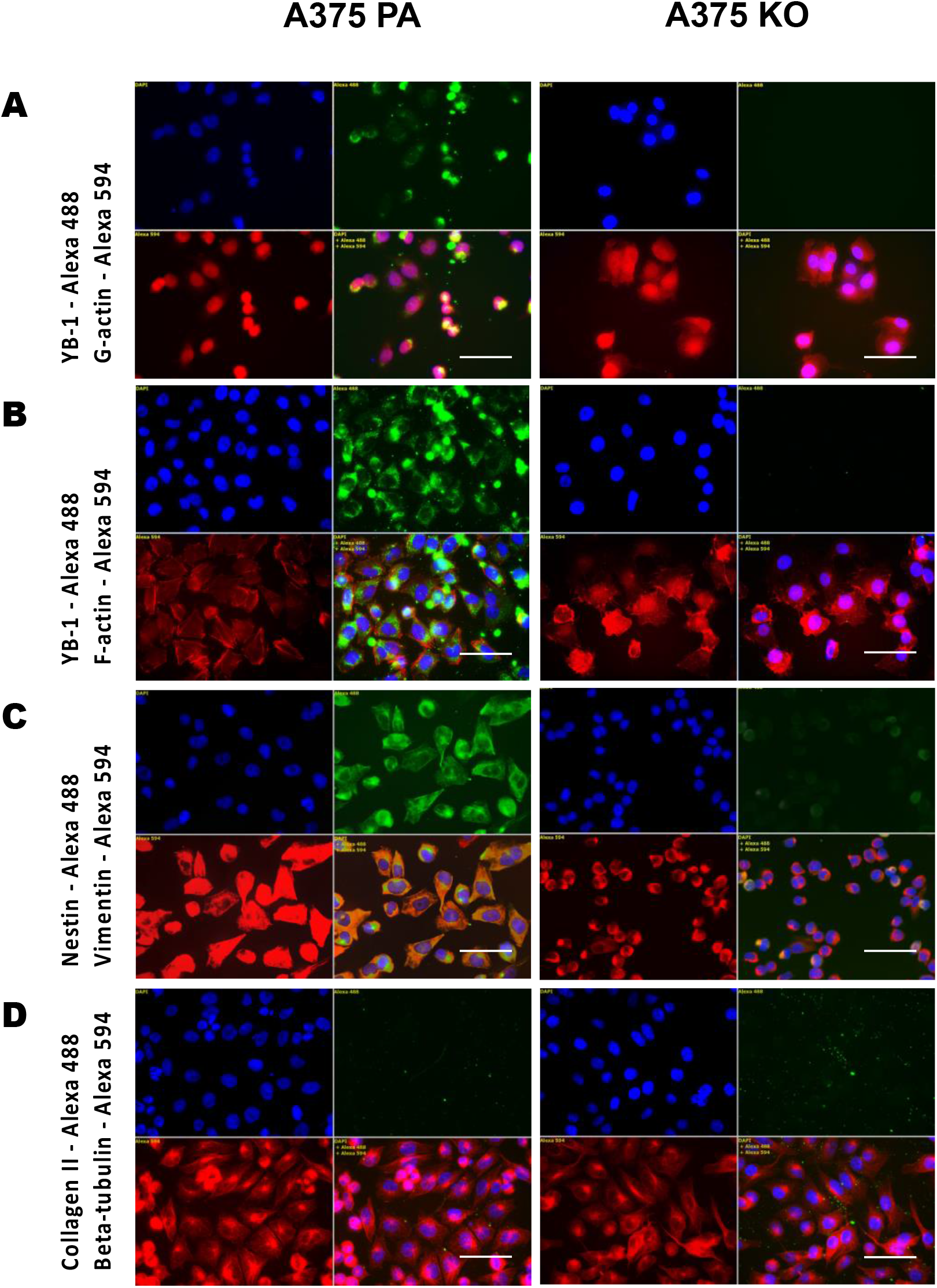
Immunolabelling of A375 parental and knock-out cell lines. Immunofluorescence staining of YB-1, F-actin, G-actin, nestin, vimentin, collagen II, beta-tubulin were conducted for these two cell lines. Alexa 488 - green labeling, Alexa 594 - red labeling, DAPI - blue (nucleus staining). Scale bar (white) represents 20 µm. Abbreviations: PA - parental; KO - knock-out.

## 4. Discussion

Malignant tumors have a high capacity to adapt to the selective pressures they encounter at every stage of tumor development, from early stages of tumor initiation, through disease progression, and in response to therapy. This plasticity is ensured by EMT (28). YB-1 has been shown to induce a highly motile, invasive mesenchymal phenotype in various cancers including melanoma (9,29,30). In the present study, we aimed to further elucidate the role played by YB-1 in melanoma cells, the biomechanical properties related to the biochemical composition of the cytoskeleton, and its rearrangement as well as upregulated proteins. To this end, we employed both quantitative as well as qualitative and biomechanical (AFM) and biochemical (proteome analysis and immunolabelling) techniques.

The general consensus on the role of the cytoskeleton in the change of metastatic potential is that a reduction in actin filaments, but an increase in tubulin allows cells to increase their invasiveness. Previous studies showed that melanoma cells with the higher Young modulus (higher stiffness) were less capable to penetrate different barriers than cells with the lower Young modulus (lower stiffness) (31). While the increase in cell elasticity correlates with progression from normal to transformed malignant cells, migrating cells generally have thicker stress fibers than non-motile cells which results in a higher value of Young’s modulus (32,33). Our AFM results showed that YB-1 expressing cell lines were significantly stiffer than YB-1 knock-out. Immunofluorescence staining revealed a less rich actin network and augmented tubulin expression in YB-1 expressing cell line compared to YB-1 knock-out. Nestin has been recently considered as a marker of malignancies of neuroectodermal origin including melanoma, where it was recognized as a prognostic factor (34). Nestin may also play a role in connecting components of the cytoskeleton as it copolymerizes into filaments with class III IF proteins, mostly vimentin, contributing to its disassembly. Interestingly, concomitant expression of YB-1 was also associated with significant preferential expression of vimentin, which is widely expressed in mesenchymal cells and has been recognized as a marker of EMT where immotile cells convert to motile mesenchymal cells (35). In previous studies, vimentin has been shown to enhance cellular elasticity (36), which was not observed in our study. Vimentin and nestin are considered to be important and specific markers for highly invasive melanoma (34). In contrast, in the knock-out cell line, nestin was not expressed which was also confirmed by the proteomic analysis and corroborated by the immunolabelling. We also employed proteome and further *in-silico* analyses to further understand the landscape of upregulated proteins, molecular function, and biological processes associated with YB-1 expression. The most upregulated protein in the parental cell line, following YB-1, was nestin, a class of VI intermediate filament protein. Among significantly upregulated proteins in the parental cell line was also vimentin, corroborated also by our immunolabelling results.

We also found that Fascin-1 (FSCN1) was another significantly upregulated protein belonging to the cytoskeleton. Fascin-1 is a filamentous actin-binding protein that crosslinks actin microfilaments into tight bundles. These are important for the formation of microspikes, filopodia, and invadopodia and their functionality in cell migration, cell-matrix adhesion, and cell-to-cell interactions (37). Another upregulated protein was septin-9 (SEPT9), which is the only member of the septin family that directly promotes actin polymerization and cross-linking and is involved in SEPT9-dependent F-actin cross-linking which enables the generation of F-actin bundles required for the sustained stabilization of highly contractile actomyosin structures (38). This regulatory axis of actomyosin function in melanoma is required for invasion and metastasis (39). YB-1 presence might actually induce the highly invasive phenotype with the rounded-amoeboid cell migration phenotype with distinct elastic features, as observed also in our AFM results. Our AFM and immunohistochemical results revealed perinuclear redistribution of nestin in the parental (YB-1 expressing) cell line which probably contributed to the significantly higher stiffness but did not necessarily correlate with lower invasiveness. This phenotype involves rapid changes in a migratory mode that arise as a reply to the current specifics of the environment (40). Different cell morphology of melanoma cell lines was also observed under the AFM microscope; the parental cell line exhibited round features with a larger overall size while the knock-out cells were smaller and more elongated (Figure 1). Enrichment analysis of a YB-1 expressing melanoma cell line revealed that overrepresented biological processes were strongly associated with metabolic processes, ribosomal turnover, and regulation of monocytes and leukocytes. Interestingly, the biological processes enriched in this highly invasive phenotype revealed negative leukocyte regulation and negative regulation of hematopoiesis. In melanoma, monocytes may contribute to tumor progression in part by mediating (41) tumor-induced immunosuppression while a low monocyte-to-lymphocyte ratio is characteristic of an aggressive (42) phenotype. Moreover, both vimentin and dectin-1 (DCT1) were significantly upregulated in the parental cell line and both proteins have been shown to have the ability to activate monocytes. Differential regulation of monocytes behavior is a concomitant process in the invasive phenotype.

Interestingly, overrepresented biological processes in a YB-1 knock-out cell line were strongly associated with glucosamine catabolic, metabolic, and biosynthetic processes, as well as protein tyrosine regulation and protein phosphorylation. Overall, this suggests a strong cellular pathway activation and modulation related to cell surface proteoglycans (PGs) which consist of N-acetylglucosamine and glycosaminoglycans (GAG). PGs modulate cell adhesion and motility, while GAGs play a role in the modulation of integrin function. Metastatic melanoma has been shown to increase in both glycolysis and oxidative phosphorylation (43), giving the cancer cells a metabolic advantage during the invasion of various tissues. One of the top enriched biological processes in the YB-1 knock-out cell line was the biosynthesis and metabolic process of UDP-N-acetylglucosamine (UDP-GlcNAc). Aberrant glycosylation is a well-established hallmark of cancer; in melanoma, N-glycosylation (44) contributes to melanoma metastasis and progression. In previous studies, nestin depletion influenced signaling through integrin and PI3K/AKT/mTOR pathways (45) and led to increased focal adhesion kinase abundance and phosphorylation.

Noteworthy to mention is that the landscape of upregulated proteins in the absence of YB-1 included zyxin which is a phosphoprotein implicated in actin cytoskeleton assembly and is mainly localized at focal adhesions which link the ECM to the actin cytoskeleton. Zyxin expression was found to be directly related to cell spreading and proliferation and inversely related to differentiation (46). Strong focal adhesions (FA) are characteristic of the elongated mesenchymal invasive phenotype. Both N-glycosylation and zyxin upregulation (47,48) would suggest that YB-1 knock-out could also be prone to metastasize. Other strongly upregulated proteins included TGM2 which is a stress-responsive gene that encodes a multifunctional and structurally complex protein called tissue transglutaminase (TG2). TG2 has been associated with the development of chemoresistance in malignant melanoma cells by exploiting integrin-mediated cell (49) survival signaling pathways. 14-3-3 protein sigma is an adapter protein implicated in the regulation of a large spectrum of both general and specialized signaling pathways, when bound to KRT17, stimulates Akt/mTOR pathway and indirectly activates p53/TP53 (50). There is increasing evidence that inactivation of 14-3-3σ is involved in tumor development in a variety of malignant tumors, therefore its strong expression in the YB-1 knock-out could be triggered by lower tumorigenicity or perhaps loss of inhibition otherwise activated in the presence of YB-1.

Collectively, constitutive expression of YB-1 seems to regulate complex and intrinsic networks and cellular pathways which overall result in amoeboid migration. This phenotype is stiffer and, in agreement with previous studies, regulates immune cell functions. Concomitant YB-1 expression also results in a strong expression of nestin and vimentin networks, as well as fascin and septin-9 which is important for the stabilization of actomyosin structures. YB-1 knock-out resulted in complete nestin depletion and strong upregulation of glucosamine catabolic, metabolic, and biosynthetic processes modulating focal adhesions. Strongly enriched N-glycosylation and zyxin biosynthesis would suggest that YB-1 knock-out acquires some characteristics of mesenchymal phenotype but lacks important markers of malignancy and invasiveness such as nestin or vimentin. We posit that there is an association of YB-1 expression with an amoeboid phenotype, which would explain the increased migratory capacity.

## Conclusion

Despite advanced treatment strategies, such as targeted therapy options and immunotherapies (immune checkpoint blockade), survival rates of humans diagnosed with advanced melanoma remain poor. Expression of YB-1 regulates complex and intrinsic networks and cellular pathways which overall render cells stiffer and acquire the ameboid phenotype which explains the increased migratory capacity.

## Acknowledgments

We would like to thank the Proteome Center, University of Tübingen, Germany, for conducting the NanoLC-MS/MS data generation.

## Authors’ contribution

AC performed the experiments, did the statistical analyses, and wrote the manuscript; UKH designed the study, supervised the work, and critically revised the manuscript; AT designed the study; CK provided the cell lines and helped to interpret the data; RR helped with the experiments and to interpret the data; MD designed the study, supervised the work, and co-wrote the manuscript. All authors critically read and approved the final manuscript.

