## Supplementary Figure 1 for "Biomechanical and biochemical assessment of YB-1 expression in melanoma cells"

### Supplementary Figure legends

A

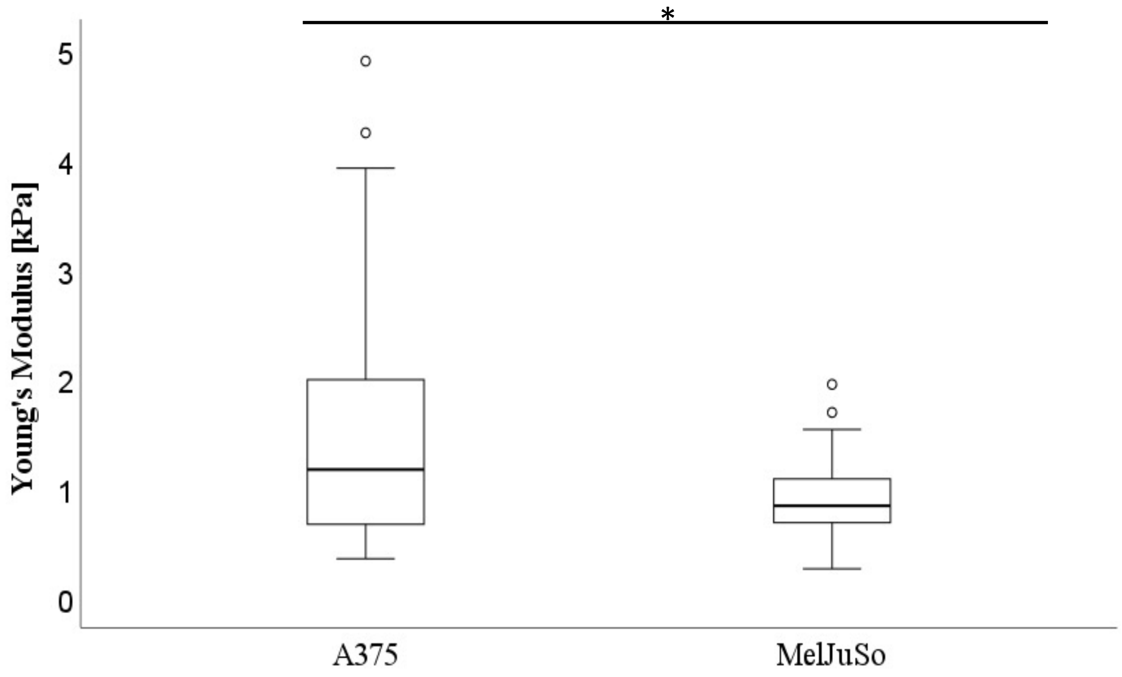


B

| Descriptive statistics | A375 [kPa] | MelJuso [kPa] |
|---|---|---|
| Median | 1.16 | 0.84 |
| Minimum | 0.35 | 0.26 |
| Maximum | 4.90 | 3.14 |
| Mean | 1.52 | 0.97 |
| Standard deviation | 1.07 | 0.55 |
| Standard error | 0.11 | 0.50 |

**Supplementary Figure 1. Cell stiffness as measured by the Young's modulus (kPa) is higher in the YB-1 expressing parental A375 cells when compared to the MelJuso cell line**. Boxplots represent the difference in stiffness (kPa) measured by AFM between the two cell lines (A375 and MelJuso) (**A**), with the descriptive statistical analysis below (**B**). A significantly higher stiffness was observed in the A375 cell line compared to its corresponding MelJuso cell line ($p < 0.001$). *$p < 0.05$; Abbreviations: PA - parental; KO - knock-out; AFM - atomic force microscopy.
